# A Pan-Cancer assessment of alterations of the kinase domain of ULK1, an upstream regulator of autophagy

**DOI:** 10.1101/702522

**Authors:** Mukesh Kumar, Elena Papaleo

## Abstract

Autophagy is a key clearance process to recycle damaged cellular components. One important upstream regulator of autophagy is the ULK1 kinase. Several structures of the ULK1 kinase domain have been solved, but a comprehensive study, including molecular dynamics, is missing. Also, an exhaustive description of ULK1 alterations found in cancer samples is presently lacking. We here applied a framework which links -omics data to structural protein ensembles to study ULK1 alterations from genomics data available for more that 30 cancer types. Moreover, we predicted the effects of mutations on ULK1 function and structural stability, accounting for protein dynamics and the different layers of changes that a mutation can induce in a protein at the functional and structural level.

We find that ULK1 is down-regulated in gynecological tumors. In other cancer types, ULK2 could compensate for ULK1 downregulation and, in the majority of the cases, no marked changes in expression have been found. 36 missense mutations of ULK1 are co-occurring with mutations in a large number of ULK1 interactors, suggesting a pronounced effect of the upstream steps of autophagy in many cancer types. Moreover, our results pinpoint that more than 50% of the ULK1 mutations that we studied are predicted to affect protein stability. Three mutations (S184F, D102N, and A28V) are predicted with only impact on kinase activity, either modifying the functional dynamics or the capability to exert effects from distal site to the functional and catalytic regions. The framework here applied could be extended to other protein targets to aid the classification of missense mutations from cancer genomics studies, as well as to prioritize variants for experimental validation, or to select the appropriate biological readouts for experiments.

## INTRODUCTION

Autophagy is a highly conserved catabolic mechanism across eukaryotes to degrade different cellular components and molecules, including organelles, proteins and bacteria ^1,2^. Autophagy initiates with the formation of the autophagosome, which later fuses to the lysosome, resulting in the degradation of the cargo and the release of cellular building blocks ^3,4^. At a basal level, autophagy contributes to maintain cellular homeostasis and the recycling of cellular components. Autophagy can also be induced as a response to different stresses with a cytoprotective role. Defects in the autophagy machinery are often linked to diseases, including cancer, neurodegeneration and bacterial infections ^5^.

Autophagy is initiated through the ULK1 (Unc-51 like autophagy activating kinase 1) complex ^6–8^, which consists of the ULK1 protein kinase, the FIP200 (FAK family kinase interacting protein of 200 kDA) scaffold protein, and ATG13-ATG101 HORMA (Hop/Rev7/Mad2) complex ^9^. The ULK1 complex can integrate different signals to promote both bulk and selective autophagy ^9^. A highly coordinated and conserved cascade of post-translational events, including phosphorylation and ubiquitination, of the ULK1 kinase complex acts as major switches for autophagy initiation ^10–13^. In the majority of the cases, the autophagy initiation is regulated by the interplay between mTOR (mammalian Target of Rapamycin) and AMPK (AMP-activated protein kinase) kinase complexes, which perform a series of inhibitory or activatory phosphorylations of the ULK1 kinase complex in different physiological conditions ^10,14–18^. Upon activation ULK1 directly phosphorylates the PI3K kinase complex including BECLIN-1, VPS34, and ATG14, which facilitate nucleation of phagophore membrane at the phagophore assembly sites ^19–21^. ULK1 also phosphorylated AMBRA1, which is a positive regulator of the PI3K complex ^21^. By analogy to the yeast counterpart, ULK1 has been suggested to play essential scaffolding roles for autophagosome formation and maturation ^9^. Mammals have five ULK1 homologs with ULK1 and ULK2 featuring the highest similarities and functional redundancy ^7^, implying that both need to be inactivated for a substantial inhibition of autophagy ^22^. ULK1 has been reported as overexpressed or downregulated in different cancer types or subtypes ^23–26^. Autophagy, in general, has a strong association with cancer and can act with a dual role in a context-dependent way, being both tumor suppressor or promoter ^27,28^. AMPK-ULK1 mediated autophagy induces resistance against bromodomain and extraterminal domain inhibitors, which are novel epigenetic therapeutics for acute myeloid leukemia ^29^. Several inhibitors of ULK1 have been used to study its function in autophagy ^21,30–32^. These molecules have potential to be used in cancer therapy since cancer addiction to autophagy has been reported ^33^. Moreover, ULK1 is involved in the first biochemical steps of autophagy, representing an amenable druggable target in the pathway. X-ray structures of ULK1 in complex with inhibitors are available (PDB entries: 4WNO ^31^, 4WNP ^31^, 5CI7 ^32^, 6QAS ^34^, and 6MNH ^35^).

ULK1 is a multi-domain serine-threonine kinase, enriched in disordered regions ^36^, which is a common trait of many scaffolding proteins. ULK1 has preference for serine as the phospho-acceptor residue in the substrate and for hydrophobic residues surrounding the phosphorylation site ^21^. The kinase domain is located at the N-terminal region of the protein and conserved among yeast and mammals ^9^. The N-terminal kinase domain of ULK1 includes the activation loops (165-174 and 178-191) and the catalytic loop (136-145), which are required for its function. ULK1 also includes a proline/serine-rich region (279-828) and a C-terminal domain (829-1051), which are involved in interactions with ATG13, FIP200 and other upstream regulators such as mTOR and AMPK ^15,37^. On the contrary, the N-terminal kinase domain of ULK1 could interact with the LRKK2 protein ^38^. An important activatory autophosphorylation site is located in the activation loop at T180 ^9^ which, upon phosphorylation can engage in salt bridges with the neighbouring arginine residues R137 and R170 ^31^.

Despite the large number of X-ray structures of the ULK1 kinase domain, no extensive molecular dynamics (MD) simulation studies of the protein have been undertaken. The availability of a conformational ensemble of a protein is of paramount importance to better understand its function and the effects of its alterations due to mutations ^39–42^. We thus used all-atom and coarse-grain models to obtain an ensemble of conformations and account for ULK1 flexibility and dynamics. We then applied methods inspired by network theory to achieve a Protein Structure Network representation ^43,44^ of the conformational ensemble. This methodology can help in the identification of residues important for structural stability and function, along with to pinpoint possible and elusive effects triggered by distal sites with respect to the functional residues of the protein ^45–47^.

We combined these structural studies with a curation of different layers of alteration of ULK1 in more than 30 cancer studies available in The Cancer Genome Atlas (TCGA) ^48,49^, so that changes in expression level or due to mutational events of the protein itself or its interactome can be both addressed to clarify the atlas of alteration of the protein in different cancer types. Indeed, TCGA is among the most important examples of large-scale cancer genomics studies, collecting clinical and molecular data for over 33 tumor types and more than 11000 samples from cancer patients. In addition, since the pool of normal samples is often underrepresented or not available in TCGA, the Recount and Recount2 initiatives ^50,51^ aimed at integrating TCGA data with normal healthy samples for the Genotype-Tissue-Expression (GTEx) project ^52^. The combination of analysis of -omics data and structural methods used in this study, expanding a framework that we recently applied to other proteins ^53,54^, provide a detailed assessment of the different effects that ULK1 mutations can cause to perturb the protein structural stability or activity. Our results can guide the selection of ULK1 as a target in certain cancer types, suggest which readouts to study for experimental research in cancer cellular biology, and provide knowledge for pharmacological or clinical-oriented efforts.

## RESULTS AND DISCUSSION

### Alterations in gene expression in ULK1 and ULK2 in different cancer types

In our study, we focus on annotating the effects of mutations found in cancer samples in the ULK1 kinase domain, since it is the only region with an available experimental structure, whereas the majority of the rest of the protein is predicted to be disordered according to analysis with *FELLS* ^55^. Nevertheless, due the complexity of alterations occurring in tumors, it is also important to verify whether the expression levels of ULK1 are altered and could (partially) compensate for the effect induced by the mutations. Thus, we analyzed the changes in gene expression of ULK1 using RNASeq data from TCGA ^49,56^ and Recount2 ^51,57^ initiatives, for a total of 30 different cancer types (Figure 1, **Table S1**). Due to the high functional redundancy of ULK1 and ULK2 ^58^, we also monitored the changes of ULK2 in the same cancer types. We used differential expression analyses to estimate the changes in expression of all the genes in the dataset and then retrieved the estimate for ULK1 and ULK2 (Figure 1). We observe different changes depending on the cancer types. Brain, gynecological and esophagus cancer types feature a downregulation of both genes, suggesting an impairment of their function in autophagy. In another group of tumors, compensatory effects are in play with one of the two ULK genes downregulated and the other upregulated, as exemplified by the Lung Squamous Cell Carcinoma, LUSC (Figure 1). In other cases, only one of the two kinases is up- or downregulated, suggesting that the unaltered levels of the remaining kinase can have a partial compensatory effect. No marked changes in expression of either ULK1 or ULK2 have been identified in eleven cancer types used in our study, such as breast, bladder, and certain kidney subtypes.

**Figure 1.**
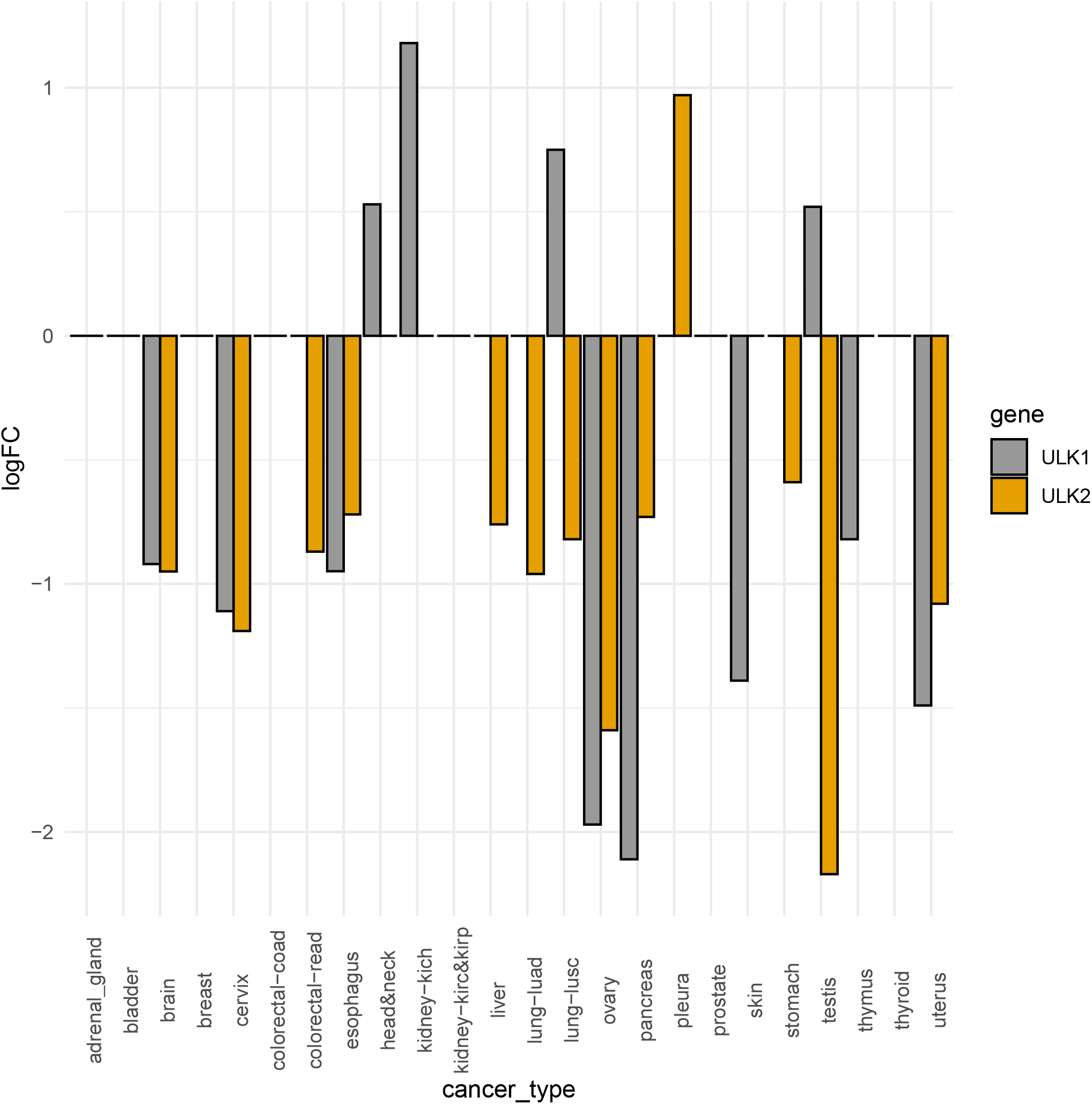
Changes in gene expression for ULK1 and ULK2 in the cancer datasets under investigation. In the plot, we report the Log Fold Change (logFC) comparing tumor versus normal samples, as estimated by *limma-voom* and including in the design matrix information on the Tissue Source Site (TSS). If the logFC > 0.05 the gene is considered up-regulated in the tumor with respect to the normal samples, whereas if the logFC is <-0.05, the gene is down-regulated. When the logFC is set to 0 in the plot, it means that the gene is not differentially expressed according to Differential Expression Analysis (see Materials and Methods for more details). The full list of logFC values is reported, together with the sample size and detailed information on the cancer datasets in Table S1.

### Functional elements of ULK1 kinase domain of interest for the study

ULK1 kinase domain (residues 8-280) consist of catalytic and regulatory regions and has been grouped under the CAMK (Ca2+/calmodulin-dependent kinase) family based on a structurally validated alignment of the kinome ^59^. It can be divided in to a small N-lobe (residues 8-92), a hinge region (93-EYCNGG-98) and a large C-lobe (99-280), as shown in Figure 2A. Before illustrating the results of our analysis, we aim at orienting the reader on the important functional ULK1 elements that we will recall in the discussion of the results. The structural background illustrated below is needed to be able to appreciate the effects that the mutations found in cancer samples could exert on this protein.

**Figure 2.**
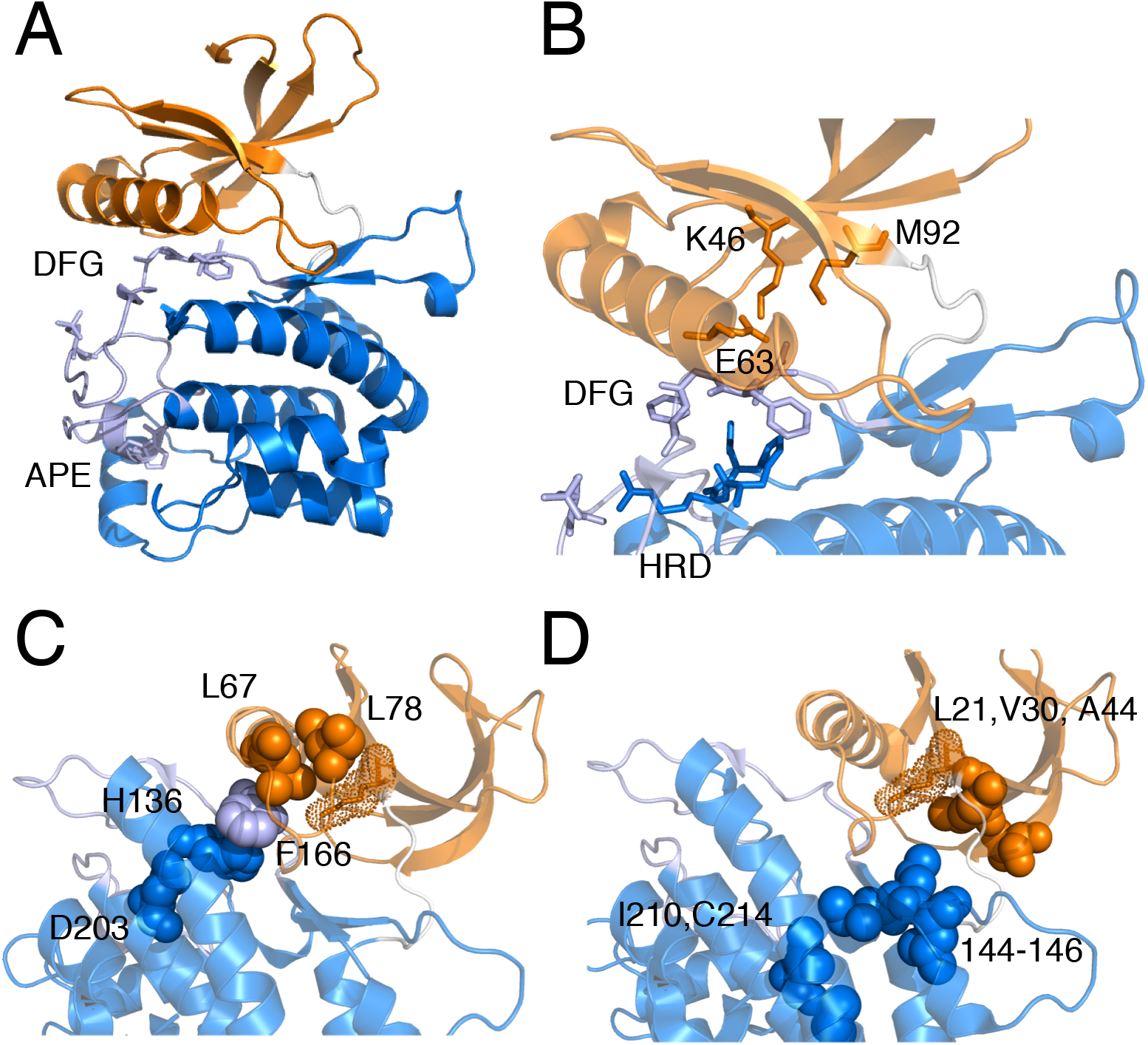
Structural and functional element of the ULK1 kinase domain, described by similarity with the c-SRC kinase. We highlight on the structure of the ULK1 kinase domain (PDB entry 4WNO) important elements of kinase function. **A)** The N-(residues 8-92) and C-(residues 99-280) lobes are shown, along with the hinge region (93-EYCNGG-98) of ULK1. The region of the activation loop (163-200) is highlighted in light blue, along with a stick representation for the side chains of the phosphorylation site T180, the DFG and APE motifs. **B)** The residues important for catalysis and substrate binding are shown. In particular, the catalytic lysine (K46) and its salt bridge with E63 in the N-lobe are shown as orange sticks, along with the gatekeeper methionine (M92). The salt bridge is also maintained in all the structures sampled by the MD simulations collected in this study (see GitHub repository). F168 adjacent to the DFG motif, which is likely to be important for substrate specificity, along with the HDR motif are also highlighted as sticks. The residues of the aligned regulatory spine (**C**) and of the catalytic spine (**D**) are shown as spheres. The catalytic lysine with dots and sticks as a reference of the active site.

ULK1 kinase domain includes a long positively charged activation loop (165-174 and 178-191) that may play a role for substrate recognition and activity regulation (Figure 2A). ULK kinases share the long disordered activation loop, a feature which is rare in the rest of the kinome ^31^. The kinase domain can be activated by phosphorylation on Thr180 on the activation loop ^9,60^ Figure 2A). The activation loop of ULK1 also includes the invariant kinase DFG motif (Asp165-Phe166-Gly167) and extends up to the APE motif (Ala189-Pro190-Glu191). In active kinases, the activation loop generally forms a cleft for substrate binding. The bound substrates form specific interactions with a conserved HRD motif (His136-Arg137-Asp138 in ULK1) (Figure 2B). The active state generally exhibits a salt bridge between a conserved lysine in the β3 strand (K46 in ULK1) and a glutamate residue (E63) in the C-helix (48-69, Figure 2B). This salt bridge is conserved in the structures from the MD ensembles collected in this study (see below). A basic patch has been observed in ULK1 around K162, which is also an acetylation site important for protein activation ^61^. This basic patch can be involved in interactions of ULK1 with membrane structures or ATG13, along with for the binding to its own C-terminal domain ^31^.

ULK1 is known to prefer serine in the target substrates ^62^, a trait that we predict related to the presence of the F168 as DFG+1 residue (Figure 2B), in agreement with the findings by Chen et al. ^63^ that large hydrophobic DFG+1 residues promote Ser phosphorylation.

Kinases often present a gatekeeper residue in the active site ^64^, which corresponds to Met92 in ULK1 (Figure 2B). Mutation of gatekeeper residues in other kinases has been associated with the development of chemotherapeutic resistance ^65^. Mutations in the gatekeeper position of kinases have been also exploited to implement a strategy for improving inhibitors potency, favouring large-to-small mutations at this site ^66^. In this context, methionine (as observed in ULK1) is among the larger and bulky gatekeeper residues found so far in kinases. Mutations of a threonine gatekeeper to methionine was associated to the development of drug resistance in other kinases ^67^.

A glycine-rich loop in the proximity of the DFG motif (G25 and G23 in ULK1) is also important for kinase function ^68^.

The regulatory (RS0: D203, RS1: H136, RS2: F166, RS3: L67 and RS4: L78, Figure 2C) and the catalytic spines (L21, V30, A44, L145, I144, L146, I210 and C214, Figure 2D) observed for other kinases are conserved in ULK1. A comparison of active and inactive regulatory spines (R-spines, RS) of kinases showed that the RS3 residue in the C-helix of the dormant enzyme is displaced and the spine not properly aligned when compared to the active enzyme ^69^. The R-spine consists of residues from both the N- and C-lobes. The histidine of the HRD motif and the phenylalanine of the DFG motif also contribute to the R-spine formation. The V30 and A44 of the catalytic spine of ULK1 should be the residues for the binding with the adenine group of ATP, whereas one of the leucine residues is likely to be important for the interaction with the adenine base. The catalytic and the regulatory spines control catalysis by dictating the positioning of the ATP and the substrate, respectively. Thus, their proper alignment is necessary for the assembly of an active kinase.

### ULK1 microsecond dynamics

Kinases are characterized by very complex conformational changes and several dynamic elements ^70,71^ and biomolecular simulations proved their effectiveness in study structure-function-dynamics relationships in kinases ^70^. We collected all-atom Molecular Dynamics (MD) simulations in explicit solvent to provide the first description of the ensemble of conformations of the ULK1 kinase domain in solution. An ensemble of conformations, as the one provided by MD, is also pivotal to the structural analyses required for the annotation of the ULK1 mutations found in cancer genomics studies, as we recently applied to other target proteins ^46,54^. To rule out dynamic patterns that are depending from the physical model used in the simulations, we collected MD simulations of the ULK1 kinase domain with two different force fields (i.e., CHARMM22star and CHARMM27). Using a dimensionality reduction approach based on Principal Component Analysis, we compared the conformational sampling ^72^ achieved with the two force fields (Figure 3A). We find a good overlap in the subspace described by the two first principal components, which account alone for more than 40% of the atomic fluctuations, suggesting that the two simulations give a consistent view on the ULK1 dynamics. To quantify the overlap between the conformational space sampled by the two different force fields, we also calculated the root mean square inner product (RMSIP), which is a measure of the similarity of the structural space described by the first 20 principal components with a value of unity as an indicator of identical subspaces. We obtained a RMSIP value of 0.76, indicating a high similarity of sampled conformational subspaces, for the two simulations of the ULK1 kinase domain, confirming the results from inspection of the 2D projections.

**Figure 3.**
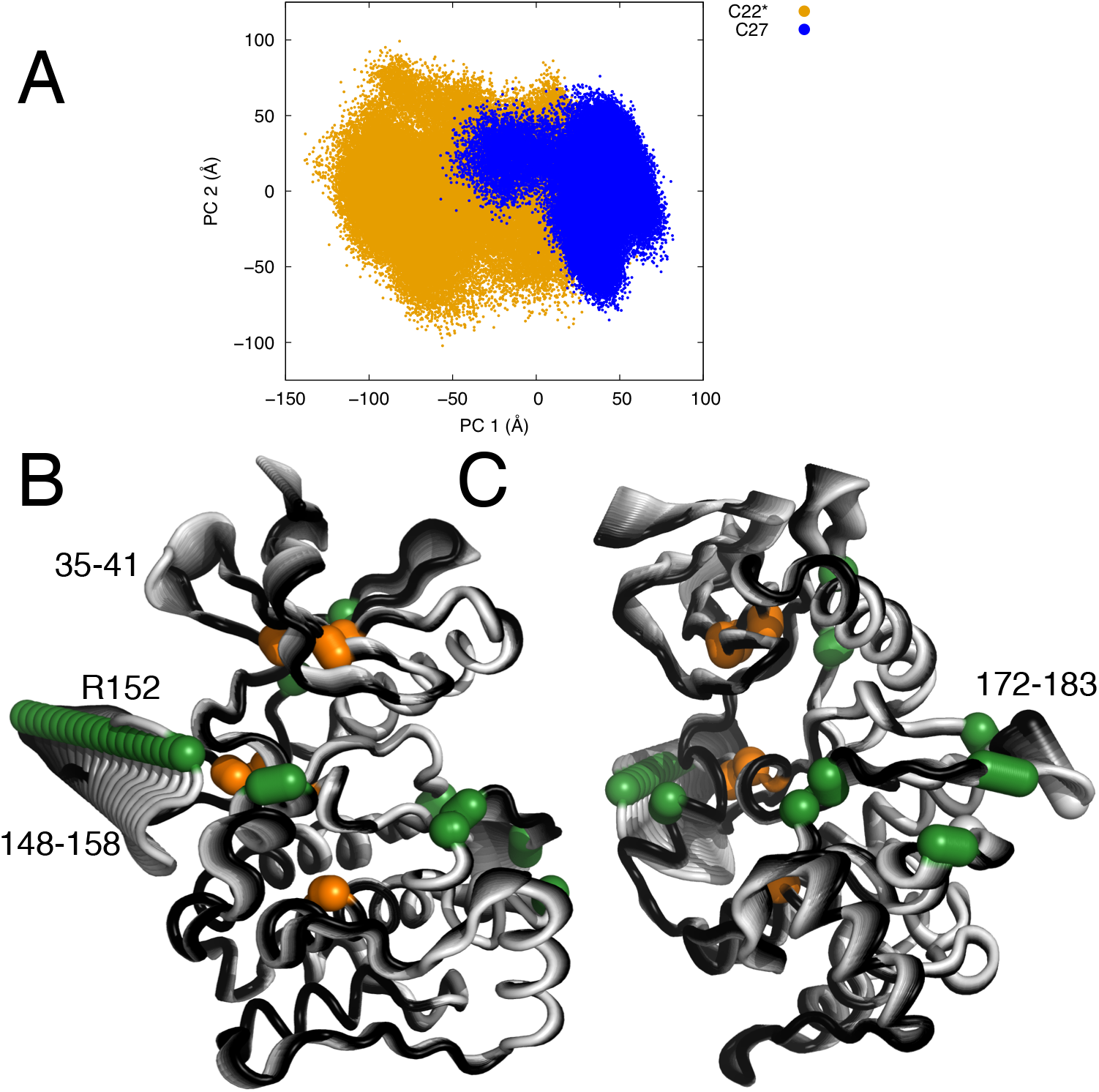
Principal component analysis of MD simulations of ULK1 kinase domain. **A)** The overlap between the sampling described by the two simulations of ULK1 carried out with two different force fields is shown along the two first principal components as a visual reference. The two MD simulations show a certain degree of overlap and sampled similar regions of subspace described by the first two principal components (PC1 and 2), with some deviations as often encountered in comparisons of MD simulations of the same system with different force fields or even different replicates of the same system. **B-C)** Conformational changes described by the first principal component, which interest three disordered regions of the protein with concerted motions. As a reference, the mutation sites in this area (discussed in the main text) and the residues important for catalysis or the regulatory spine are shown in green and orange, respectively.

Further, we analysed the principal motions described by the first principal component (Figure 3B-C) and we notice concerted motions between the loop 148-158 and a part of the activation loop (in the region 172-183), along with the loop adjacent to the catalytic lysine (in the region 35-41). The two disordered regions 148-158 and 172-183 feature motions of closure towards the rest of the ULK1 structure. The conformational change seems to be triggered by the electrostatic interactions between two arginine residues (R152 and R153) and a negatively charged residue (D102) on the facing helix. Simultaneously with this motion, a conformational change in the region 35-41 occurs where a loop moves apart from the L78 residue of the regulatory spine. The analyses of these dynamic patterns will be used in the annotations of the possible functional impact of ULK1 missense mutations found in cancer samples, discussed in the sections below.

### Missense mutations in cancer of ULK1 kinase domain

We retrieved the missense mutations in the coding region of ULK1 gene for each of the cancer studies deposited in TCGA (**Table S2**, Table 1). We identified the majority of the mutations in uterine, lung, colon and stomach tumors (i.e., UCEC, LUSC, COAD and STAD). ULK1 is also predicted as a driver gene in some of these cancer types by *OncodriveCLUST*, a method based on positional clustering and exploiting the notion that variants in cancer-causing genes are enriched at few specific loci ^73^. Most of the cancer types with ULK1 mutations are not characterized by marked changes in ULK1 expression (Figure 1), with the exception of brain and uterine tumors, where both ULK1 and ULK2 genes are down-regulated. We should notice, however, that these TCGA cancer types are characterized by a high mutational burden, as shown, for example in a recent study ^74^. Thus, it is not surprising that a higher number of missense mutations of ULK1 are found in these tumor types.

**Table 1.** Summary of the missense mutations of ULK1 kinase domain analyzed in this study. We report the mutation identified in TCGA data and located in the kinase domain of ULK1. The cancer studies in which ULK1 is predicted as driver genes are marked with a *. The full name of the TCGA cancer studies are reported in Table S1. We show also the results of the prediction of damaging or neutral mutations with *Fathmm, Revel, Sift* and *Polyphen.* We also indicate the overlap with short interaction linear motifs (SLiMs) and post-translational modifications (PTMs). The original results of each predictor are reported in Table S2. In this table we used simplified and common labels across the predictors, i.e. damaging or neutral to allow a more straightforward comparison. We also reported as a reference the results of DEA (Figure 1 and Table S1) for ULK1 and ULK2. ‘Down’, ‘up’ and ‘no DE’ indicate that the genes are down-, up-regulated, or not differentially expressed.

| MUTATION | TCGA STUDY | FATHMM | REVEL | SIFT | POLYPHEN | CO-OC. ULK2 mutations | DEA | SLiMs/PTMs |
| --- | --- | --- | --- | --- | --- | --- | --- | --- |
| G12D | GBM | damaging | neutral | damaging | neutral | NONE | down |  |
| F14L | BLCA | damaging | neutral | damaging | neutral | NONE | no DE |  |
| A28V | GBM | neutral | neutral | damaging | damaging | NONE | down |  |
| S56Y | UCEC*; COAD* | neutral | neutral | damaging | damaging | D73A;NONE | down; no DE |  |
| L78Q | STAD* | neutral | damaging | damaging | damaging | NONE | down(ULK2) |  |
| F81L | BLCA | neutral | neutral | neutral | neutral | R408S | no DE |  |
| N86T | UCEC* | neutral | neutral | damaging | neutral | NONE | down |  |
| N96D | UCEC* | neutral | neutral | damaging | damaging | NONE | down |  |
| A101T | UCEC* | neutral | neutral | damaging | damaging | T102A | down |  |
| D102N | LGG | neutral | neutral | damaging | damaging | NONE | down |  |
| D113E | HNSC | neutral | neutral | damaging | damaging | NONE | up(ULK1) |  |
| A125T | COAD* | neutral | damaging | damaging | damaging | NONE | no DE |  |
| I135V | BLCA | neutral | neutral | neutral | damaging | NONE | no DE |  |
| R137H | STAD* | neutral | damaging | damaging | damaging | NONE | down(ULK2) |  |
| R137C | COAD* | neutral | damaging | damaging | damaging | NONE | no DE | Introduce S-nitrosylation |
| D138N | LUSC;PAA D | damaging | damaging | damaging | damaging | NONE;NONE | down;down |  |
| R152L | LUAD* | neutral | neutral | damaging | damaging | NONE | down(ULK2) |  |
| A154S | LIHC | neutral | neutral | neutral | neutral | NONE | down(ULK2) |  |
| G167A | LUSC | damaging | damaging | damaging | damaging | NONE | down |  |
| A169T | KIRP | neutral | damaging | damaging | damaging | NONE | no DE |  |
| Y171H | SKCM | neutral | neutral | neutral | damaging | R951K | down(ULK1) |  |
| M177V | STAD* | neutral | damaging | damaging | damaging | NONE | down(ULK2) |  |
| G183V | LUSC | neutral | damaging | damaging | damaging | NONE | down |  |
| S184F | HNSC | neutral | damaging | damaging | damaging | NONE | up(ULK1) |  |
| S195P | UCEC* | neutral | damaging | damaging | damaging | NONE | down | predicted phosphorylation by DNAPK or ATM kinases |
| V211I | UCEC* | neutral | neutral | neutral | neutral | NONE | down |  |
| L215P | BRCA | neutral | damaging | damaging | damaging | NONE | no DE |  |
| E246D | COAD* | neutral | neutral | damaging | damaging | R254I | no DE |  |
| P250S | COAD* | neutral | neutral | neutral | neutral | NONE | no DE | introduce phosphorylation by PKC kinase |
| R252L | LUSC | neutral | neutral | damaging | damaging | NONE | down |  |
| R252W | STAD* | neutral | neutral | damaging | damaging | NONE | down(ULK2) |  |
| R261C | STAD* | neutral | neutral | damaging | damaging | V868M | down(ULK2) |  |
| R261H | PAAD | neutral | neutral | damaging | damaging | NONE | down |  |
| D268H | OV | neutral | neutral | damaging | damaging | NONE |  |  |
| F273V | UCEC* | neutral | neutral | damaging | neutral | R303C |  |  |
| D279N | UCEC* | neutral | neutral | damaging | neutral | NONE |  | IAP-binding motif (279-283, ELM) |

In total, we collected 36 different missense mutations of ULK1 kinase domain distributed over the whole structure (Figure 4A) of which D138N of the HRD motif occurred in both pancreatic and lung cancer samples and, the R137, R252 and R261 sites were mutated to different residues in different cancer types.

**Figure 4.**
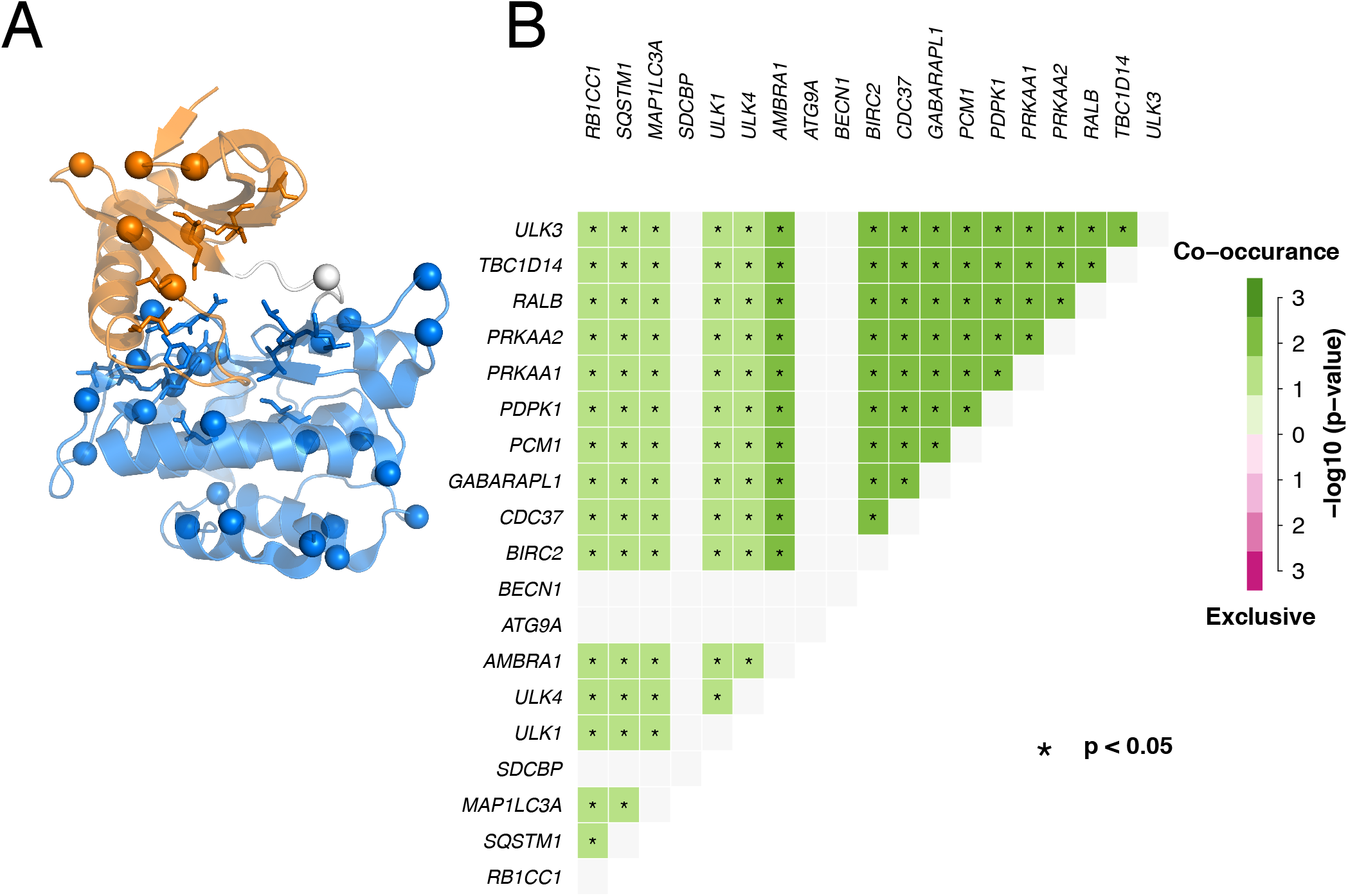
Missense mutations in the ULK1 kinase domain in TCGA datasets and co-occurrence of mutations in ULK1 interactome. **A)** The mutation sites are highlighted on the 3D structure, along with the sub-domain composition (N-, C-lobe and hinge in orange, marine and light grey, respectively). The residues of the DFG and HDR motifs are shown as sticks, whereas the catalytic lysine with sticks and dots. **B)** We used, as a description of the ULK1 interactome, the interactions annotated in *IID* and estimate the co-occurrence of mutations in the TCGA datasets under investigation. The results for the TCGA pancreatic cancer dataset are shown as an example. The other data are available in the GitHub repository associated to this publication.

ULK1 is a multi-domain protein with scaffolding functions and, as such, can interact with a multitude of different other proteins. For a proper assessment of ULK1 missense mutations, it is important to gain knowledge on the alterations, in the same tumor samples, of the biological partners of interaction.

To this scope, we curated the ULK1 interactome, mining the *IID* protein-protein interaction database ^75^ and estimated the co-occurrence of mutations among each of the 30 identified interactors and ULK1 mutations for each cancer type (**Table S2**). We found co-occurring mutations between ULK1 and its interactors in eleven cancer types in which ULK1 has been found mutated (see GitHub repository for more details on each of them). Among these, stomach, skin, brain, colorectal, uterine and pancreas tumor samples (Figure 4B as an example) are characterized by co-occurring mutations in a large number of members of the ULK1 interactome, especially the ones important for the upstream regulation of autophagy (such as AMBRA1, components of the mTOR and ULK1 complex, RB1CC1/FIP200, TBC1, AMPK subunits, members of the ATG8 family, ATG16L1, BECN-1, PDPK,I RGM, RAB1A, P62, MINK1, SDCBP, and ATG13). This suggests a pronounced effect of alterations related to ULK1 function and activity, and, more in general, upstream steps of autophagy, in these cancer types.

Methods for fast annotation of damaging and neutral mutations poorly agree on the effect of each of them (Table 1 and **S2**). Moreover, we notice that some of these methods predict several mutations with damaging effect, which is unlikely in tumor where most of the mutations will have a passenger effect, whereas others (such as Fathmm) are strict and tend to predict as damaging mutations substitutions at functional sites (such as D138N) without considering the structural impact of the mutation itself. We also verified if mutations of ULK2 kinase domain were co-occurring with mutations in ULK1 and we observed this pattern only in few isolated cases (Table 1). The mutations occurring in ULK2 kinase domain (D73A, T102A, R254I) are predicted with no effect on protein stability with the methods illustrated below, resulting in minor changes of free energy associated with stability (i.e., 0.2-1.6 kcal/mol). In the majority of the cases, we also noticed that most of the mutations occurred in cancer types where the ULK1 and ULK2 gene expression is down-regulated.

We then turned our attention to a workflow similar to the one that we recently applied to another protein ^54^ for a more comprehensive assessment and understanding of the effects induced by ULK1 mutations. We evaluated different properties to discriminate between effects associated with the structural stability of the protein or with its function. Moreover, these analyses allowed us to link the effects of the mutations with specific factors for ULK1 activity or regulation, which are important piece of knowledge to guide the experimental characterization of the molecular mechanisms and phenotypes associated with each mutation. The analyses used for the assessment are described one by one in the following sections.

### Interplay of the ULK1 mutations with post-translational modifications and functional motifs

As first factor for our assessment, we evaluated each mutation site in the context of interplay with post-translational modifications (PTMs) and overlap with functional short linear motifs (SLiMs), along with the potential of harbouring new PTM sites upon mutation (Table 1). These are properties that can ultimately influence the regulation of the protein or its spectrum of interactions.

Moreover, we evaluated if any additional mutations was found in the LC3 interaction region (LIR) of ULK1 ^76–78^, which is placed in a distal region with respect to the kinase domain. This analysis was motivated by the observation that mutations in members of the ATG8 family are co-occurring with mutations of ULK1 in some of the TCGA cancer studies (Figure 4B). LIRs are SLiMs for interaction between the LC3/GABARAP (ATG8) family members and other autophagy proteins and key mediators of autophagosome formation ^79^. We recently found mutations of LC3B co-occurring with mutations in its LIR-containing interactors in cancer genomic data ^54^ and we thus aimed here to evaluate if the same happens for ULK1. We did not find any mutations in the surrounding of the ULK1 LIR region in the TCGA samples under investigation, thus suggesting that the recognition between ULK1 and the ATG8 family is not a major driver of its alterations in these cancer samples.

The only mutation in the proximity of a SLiM is the C-terminal D279N, which includes an IAP (Inhibitor of Apoptosis Protein) binding motif (IBM). This motif has not been characterized in ULK1 yet, at the best of our knowledge. Interestingly, one of the ULK1 interactors, BIRC2 (**Table S2**) is a cellular inhibitor of apoptosis and promote autophagy, interacting with ULK1 during mitophagy ^80^. We speculate that this interaction could be mediated by the IAP motif of ULK1 and that the mutation of D279N could impair it. The mutations have been found in uterine cancer, where mutations of ULK1and BIRC2 are co-occurring (see GitHub repository).

We do not identify any mutations directly altering an experimentally validated PTM site. Nevertheless, we find one solvent-exposed mutation site (S195) which is predicted as a phosphorylation site for DNAPK or ATM kinases. Mutation to proline could have the effect to abolish this modification. We then analyzed the mutations to serine, threonine or tyrosine for their capability to introduce new phosphorylatable residues, along with mutations to cysteine for their possibility to introduce a redox-sensitive post-translational modification, i.e. *S*-nitrosylation ^81^ (Table 1, **Table S2**). P250S ULK1 variant is predicted to result in a phosphorylatable serine by PKC kinase and the R137C mutation in the HRD motif as a possible *S*-nitrosylation site.

### Assessment of the impact on ULK1 protein stability of the missense mutations

One of the main effects that a mutation can have on a protein is to alter its structural stability, causing local misfolding and a higher propensity for the mutated variant to be targeted by pathways for protein clearance, such as proteasomal degradation ^42,82,83^. In this scenario, the function of the protein will also be affected, but mostly as the result of compromised protein levels and not necessary an alteration of its capability to interact with the biological partners or be an active enzyme.

We used a high-throughput saturation mutagenesis approach ^84^ based on an effective empirical energy function implemented in *FoldX* to predict the effect on protein stability induced by all the possible substitutions of each position of the structure of the kinase domain of ULK1. This approach has the advantage of providing both the estimate of the damaging effect of the disease-related mutation of interest and a pre-computed list of predicted changes in stability for any other mutations of the protein. The latter is useful to identify important hotspots for the structure of ULK1, along with pre-annotated effects of amino acidic substitutions that can be consulted for newly discovered mutations in future genomics studies.

To overcome the inherent issues in local sampling and lack of backbone flexibility of *FoldX*, we used the MD-derived ensembles for the analysis. We estimated the changes in the free energy of folding upon mutation (**Table S3-S6**), as recently applied to other cases study ^46,54,85^. The predicted ΔΔGs, using the two different MD ensembles of the ULK1 kinase domain, are in good agreement (**Table S3**). Moreover, we notice that, for some mutations, the predicted damaging effect is a result of the usage of the static X-ray structure (**Table S3**), whereas the possibility to account for the flexibility of the protein structure in the prediction result in neutral effect. An example of this behavior is A28V (**Table S3**). Based on this observation, we used the ΔΔGs predictions from the calculation on the MD ensembles to classify the ULK1 mutations found in cancer patients (Figure 4B). Two mutations (S184F and V211I) result in a stabilization of the protein architecture, suggesting a better packing of the protein. S184 is located in the proximity of aromatic residues, including the one of the DFG motif, which is important for catalysis so it cannot be excluded that a mutation to S184F could alter the functional dynamic of the kinase at this site.

We also notice that some mutation sites are more general hotspots for ULK1 structural stability (Figure 5A). In these cases, the sites are sensitive to substitutions to most of the other residues (i.e., F14, S56, L78, F81, A125, R137, A169, G183, and F273). I135 is also a stability hotspot, but the I135V mutation, which was found in the cancer sample, is one of the few tolerated substitutions, suggesting a neutral effect.

**Figure 5.**
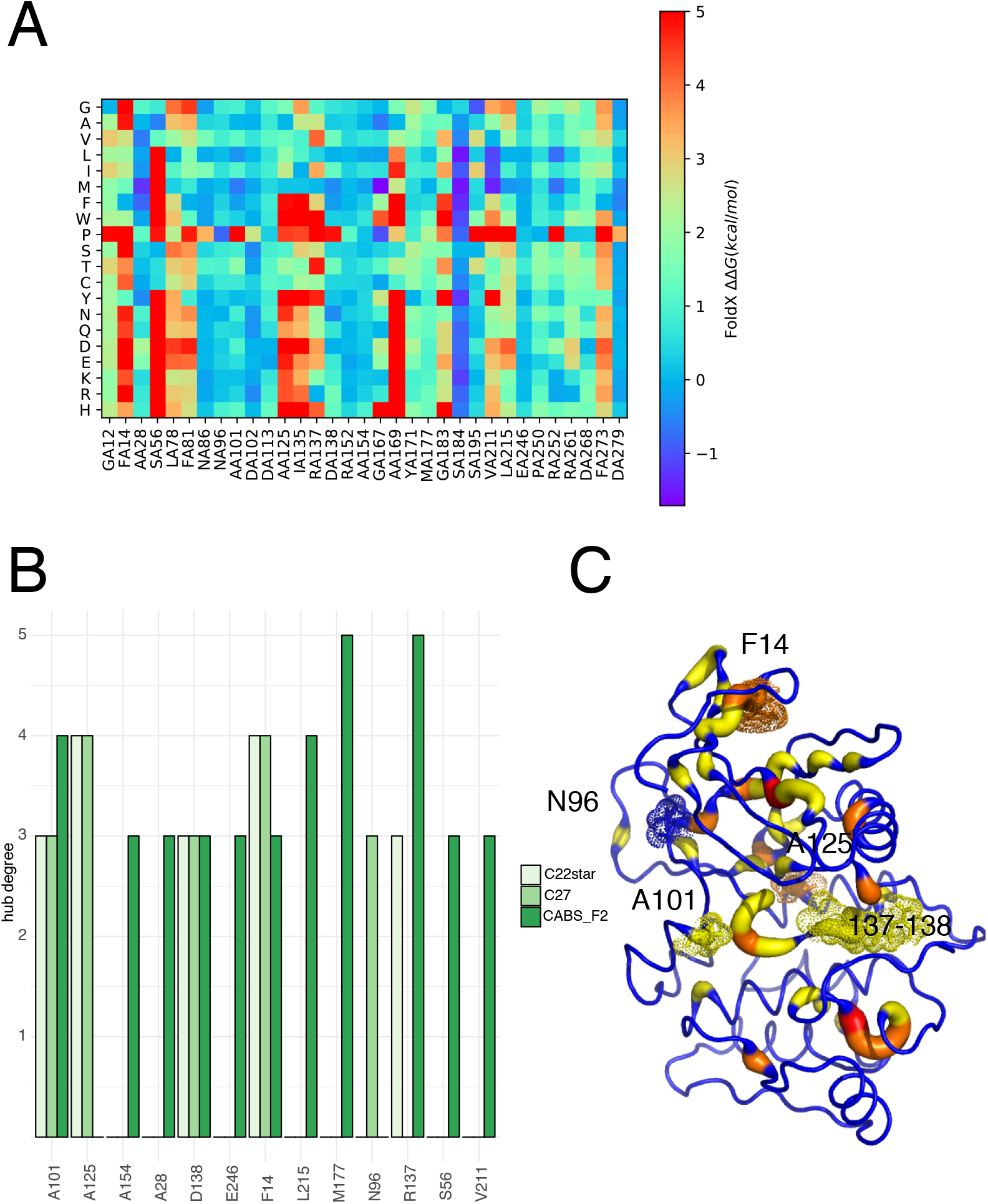
ULK1 mutations in TCGA cancer samples altering protein stability. We used two different estimates of the impact on protein stability, based on the calculations of changes in free energy of unfolding with an empirical scoring function (**A**) and the notion of hub residues in a Protein Structure Network (**B-C**). **A**) A heatmap with the estimated changes in free energy upon a deep mutational scanning of all the mutation sites is reported. As an example, we showed the results for the MD-ensemble with CHARMM22star force field. The results of the other mutational scans are reported as TableS4-S6. More than 50% of the ULK1 mutations are predicted to destabilize the protein structure and located in sites that are general hotspots for maintenance of the native fold. **B**) Hub prediction are in fair agreement using different methods to generate the ensemble of conformations, i.e. MD simulations with two different force fields (CHARMM22star and CHARMM27), along with a coarse-grained sampling method with a posteriori reconstruction of a full atom model (CABS-Flex), which overestimate some of the hubs in the C-terminal domain of the protein. **C**) Hub-behavior of the mutation site in the MD ensemble with CHARMM22star force field, as an example.

The availability of MD ensembles for ULK1 kinase domain prompted us to apply a Protein Structure Network (PSN) approach based on the persistence of side-chain contacts in the conformational ensemble ^86,87^ to estimate hub residues, which are often corresponding to important residues for protein stability (Figure 5B-C). We verified which of the mutation sites are found in correspondence or proximity of a hub, as an additional parameter to evaluate the impact of the substitution on protein stability. Moreover, to account for the fact that the mutation in a hub can still retain its hub capability, we collected the same analyses on conformational ensembles derived for each of the mutant variants (**Table S3**). We classified as damaging mutations according to this parameter only the ones for which the substitution abolishes the hub behavior. We find only one case, i.e. F14L, where the hub behaviour is conserved upon mutation. Most of the mutation sites corresponding to PSN hubs also corresponds to mutational hotspots associated with high ΔΔGs for protein stability, with the exception of D138N.

Another strategy to impair structural stability could be related to the loss of electrostatic interactions in the form of salt bridges or hydrogen bonds. We calculated the persistence of the salt bridges and their network in the MD ensembles and annotate which of the mutation site where likely to abolish these interactions (Table 2). Most of the mutations of residues involved in salt bridges are likely to have marginal effects since they conserve either the negatively charged nature of the wild-type residue or they are replaced by asparagine, which could still account for electrostatic interactions with the guanidinium group of arginine. The only mutations which could impair salt-bridge formation are R152L and D268H. R152L is involved in salt bridges in the CHARMM27 MD ensemble, whereas it shows a loose tendency to form electrostatic interactions in CHARMM22* MD ensembles. MD force fields are known to have limitations in overestimating salt bridge contributions ^88,89^ and according to a recent benchmarking ^89^, we selected the CHARMM22star results for the annotation of the mutations with respect to salt-bridge formation. Overall 58%of the ULK1 mutations found in the different cancer types are predicted damaging for protein stability using at least one of the two criteria above.

**Table 2.** Salt-bridges involving ULK1 mutation sites characterized by charged residues, along with their persistence in the MD ensemble. R137, R252 and R261 are not reported since we did not find any stable salt bridges involving these residues in the simulations.

| **MUTATION** | **CHARMM22star (%)** | **CHARMM27 (%)** |
| --- | --- | --- |
| D102N |  | R152-D102-R153(67.5, 81.0) |
| D113E | D113-R116 (44.3) | D113-R116(55.8) |
| D138N | D138-K140 (99.8) | D138-K140(99.8) |
| R152L |  | R152-D102(67.5) |
| E246D |  | E246-R245(22.7) |
| D268H | D268-K201(98.8) | D268-K201(93.0) |
| D279N | D279-R116(63.2) | D279-R116(59.4) |

### Assessment of the impact of mutations on ULK1 function

Apart from effects on the stability, the availability of structure and dynamics of ULK1 allowed us to predict effects induced by the mutations on its function.

We evaluated the occurrence of the mutations in proximity of the disordered regions interested by the functional motions underpinned by Principal Component Analysis (Figure 5B-C). R152L and D102N are likely to impair or weaken, the functional motions observed for the region 148-158, which are triggered by their electrostatic interactions. We cannot rule out that the presence of R153 in the R152L mutant variants could partially compensate for the mutation. On the other side, the lateral motion of the region 172-183 of the activation loop is surrounded by different mutation sites, such as M177 in the loop itself, S195 in the region where the loop bends, and Y171, G183/S184 which act as hinges for the motion of these regions. It could be expected that these mutations impair ULK1 functional dynamics. In addition, the regulatory spine residue L78 and F81 in the proximity of the other disordered loop (35-41), which is involved in the concerted conformational changes, are also corresponding to mutation sites.

PSN approaches can be used to infer functionally-damaging sites if the paths of communication between the mutation sites and other important functional sites for the kinase activity are considered. This analysis can shed light on effects that are likely to be transmitted long-range, often at the base of allostery ^45,47,90^. We estimated all the shortest paths of communication between each mutation site and five important classes of residues for kinase function. In particular, we selected three groups of target residues: i) residues important for activity (K46, E63, M62, T180 and K162); ii) residues of the DFG, HRD, and APE motifs; iii) central residues of the C-helix (55-65); iv) the residues of the catalytic and v) regulatory spines (Figure 6). We selected only those paths conserved both in the CHARMM22star and CHARMM27 simulations (**Table S7**). We did not find any communication roads to the regulatory spine or to the APE motif. On the contrary, a subset of mutation site (A28, A101, D102, D138, N96, and R137) was communicating with at least two of the target areas of interest, often using multiple paths. This result suggests that substitutions at these sites could be detrimental for functional long-range communication or it could increase it (if oncogenic) especially in cases in which the steric hindrance or the physico-chemical properties of the wild-type residue are not maintained upon mutation, as in the case of A28V, A101T, R137C and, R137H. We used the statistical mechanical model implemented in AlloSigMA ^91^ to have a more direct proof that these mutations could exert an allosteric effect (see Github repository for the outputs).

**Figure 6.**
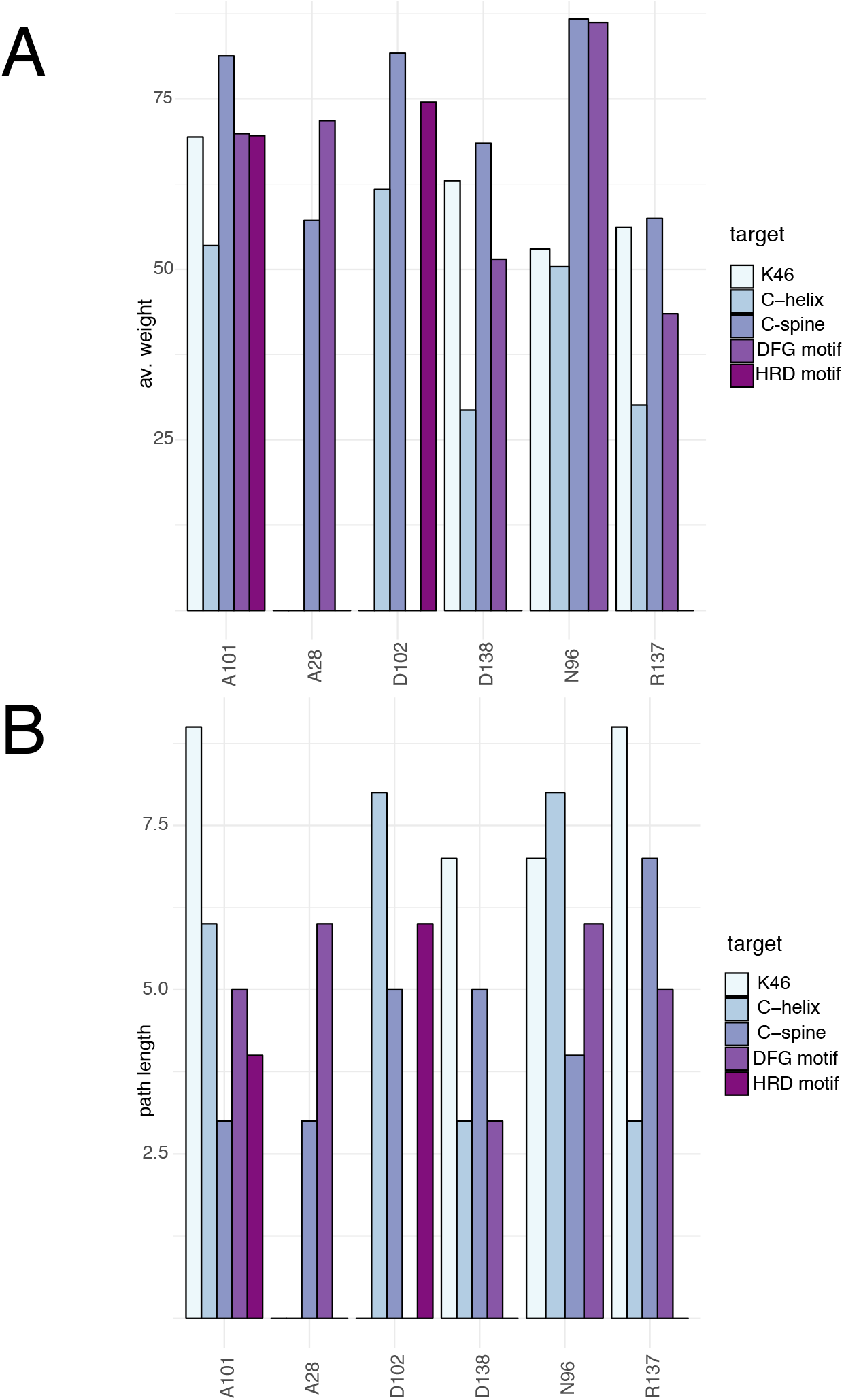
Shortest paths of communication between the mutation sites and key functional or regulatory regions of ULK1 kinase domain. The whole set of results is available in Table S7. We reported the data for CHARMM22star MD simulations for sake of clarity, which were very similar to the ones for the simulation carried out with CHARMM27. If a mutation site featured more than one path of communication for the same class of target residues, only the path with lower path length is taken into account in this figure. We used as target residues: i) residues important for activity (K46, E63, M62, T180 and K162); ii) residues of the DFG, HRD, and APE motifs; iii) central residues of the C-helix (55-65); iv) the residues of the catalytic and v) regulatory spines. No paths were found from the mutation sites to the regulatory spine or the APE motif in both the MD simulations.

### General assessment and classification of ULK1 missense mutations in TCGA

We integrated all the results collected by each of the analyses above to provide an overall view on the several properties and layers of alterations that a mutation in a protein can cause, and which are ultimately connected to the alterations in its function at the cellular level. In particular, with our framework we can assess: i) effects on protein structural stability, which will impact on the protein levels and turnover in the cell; ii) interplay with post-translational modifications and emergence of new layers of regulation; iii) alterations of binding regions for biological partners; and iv) long-range functional effects. We classified the mutations according to each of these properties as damaging or neutral (Figure 7A) and then rank them. The ranking allowed us to identify mutations that are likely to be damaging, along with to identify if the effect is triggered more by a destabilization of the protein product or a stable protein variant with impaired functionality (Figure 7B).

**Figure 7.**
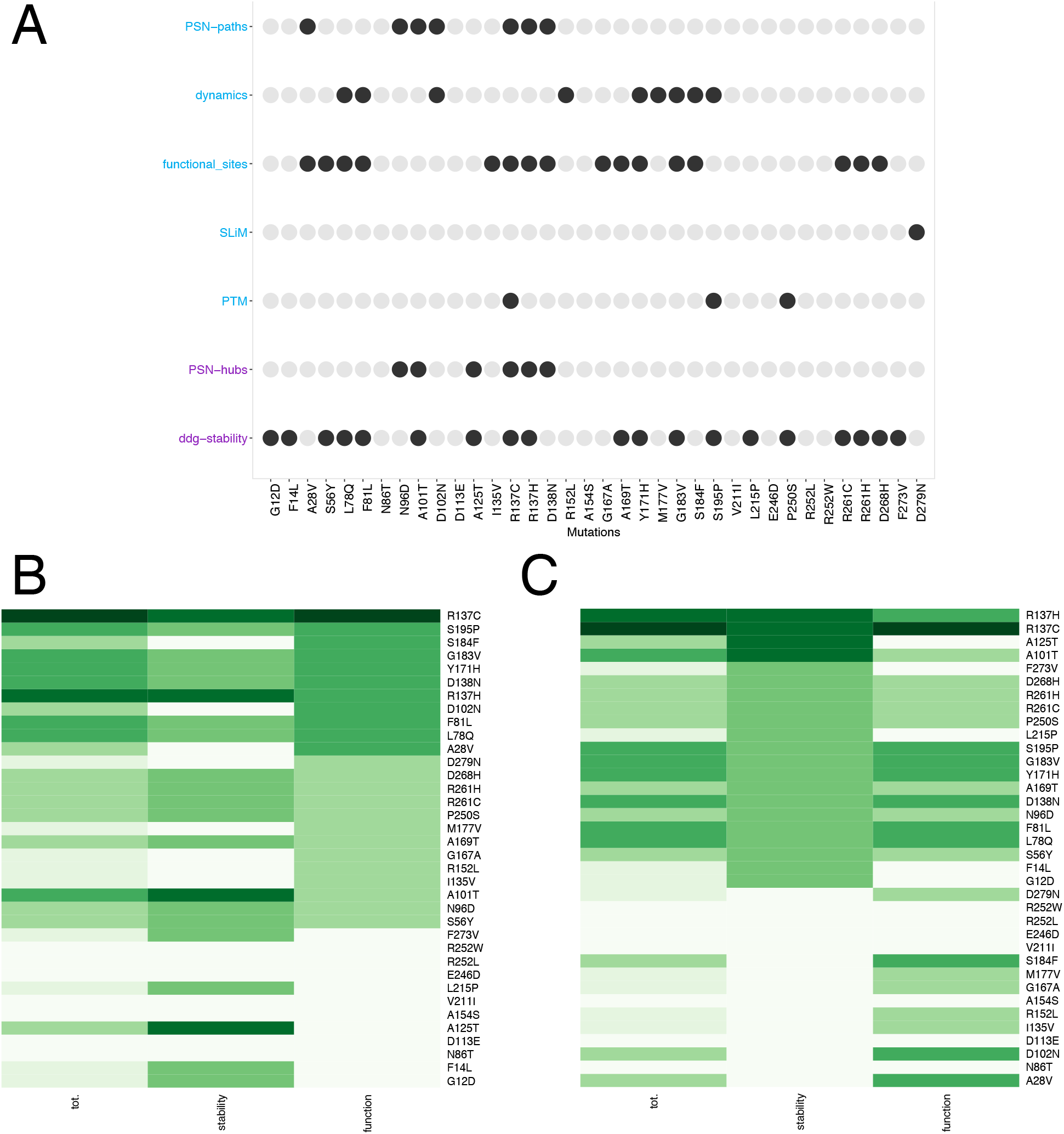
Classification of ULK1 missense mutations found in cancer genomics studies. **A)** The different analyses used in this study have been aggregated to associate the potential of damaging or neutral effect of each mutation. We used descriptors that account for either protein stability (purple) or function (blue). Mutations altering one of these properties are indicated as black dots. The diagram allows to link each mutation with a specific effect which could also guide the selection of the more appropriate set up for experimental validation. **B-C)** The heatmaps with the results of ranking on a collective score of damaging potential for the mutations in term of function (B) or stability (C). Darker the color more damaging the mutation is predicted to be.

We observed, as stated above, that more than 50% of the mutations of ULK1 found in the cancer samples are predicted to alter protein stability. This is often accompanied by a possible impact also on the native functional properties of the same variant. A minority of mutations are only damaging for stability (A125T, F273V, L215P, F14L and G12D) and do not alter the functional state of the protein. The detrimental effect on the protein stability observed by these mutations could alter the cellular level and turnover of the protein. This effect could be dominant with respect to the effects that the same mutations exert on protein activity or interactions.

ULK1 stability could also be modulated by the interaction with ATG13 and FIP200, which bind to the cytoplasmic domain of the ULK1 ^37^, or by chaperonins like proteins, such as p32 ^92^. Thus, these interactions might compensate for the loss of stability induced by some of the mutant variants. Interestingly, in several cases the cancer types where destabilizing mutations of ULK1 occurred, also feature co-occurrence of mutations in ATG13 or FIP200 (**Table S3**), suggesting an overall alteration of ULK1 stability. Three mutations (S184F, D102N, and A28V) are predicted with a possible impact only on kinase activity, either altering the functional dynamics of the protein or the capability to exert long range effects from distal site to the functional and catalytic regions.

We searched in the literature if the mutation sites under investigation have been subjects of experimental studies and if the results of this experiments corroborate our prediction. We found a study reporting a mutation of S184 to alanine ^93^. S184 is mutated to phenylalanine in head and neck TCGA samples and, we predict a marginally stabilizing effect upon mutation. Moreover, we classify S184 as a functional damaging mutation likely to impair ULK1 functional dynamics, due to its hinge behavior for the motions of the activation loop. S184A and S184D have been reported to inactivate the ULK1 kinase ^93^, supporting the predicted functional role more than an effect on structural stability. We collected other mutations at other sites for which the functional impact has been studied experimentally and they are summarized in **Table S3.** Of interest, S174A mutation results in a hyperactive enzyme ^93^ and it is located in the region of the activation loop undergoing conformational changes. Moreover, we notice that the other experimental mutations with a reported effect on ULK1 activity are predicted to have marginal effects on protein stability, with the exception of K46N, M92A and Y89A. These mutations result either in an inactivation of the kinase ^94^ or an impairment of phosphorylation of the ATG13 substrate ^25^. Our calculations suggest that the detrimental effect could come from altering the protein stability, an aspect which could deserve further investigation to verify the cellular levels and half-life of these variants to conclude which one is the predominant effect.

## MATERIALS AND METHODS

All the inputs, scripts and main outputs of this study are available in the GitHub repository https://github.com/ELELAB/ULK1_mutations. The trajectories and input files for the molecular dynamics simulations have been deposited in OSF: https://osf.io/8xuaj.

### Expression levels of ULK1 in TCGA datasets

We downloaded and pre-processed level 3 harmonized RNA-Seq data (HTSeq count) for all the available datasets from TCGA. We downloaded the data in June 2019 from the Genomic Data Common (GDC) Portal using the GDCdownload function of *TCGAbiolinks* ^95^. An overview of the analysed datasets is reported in **Table S1**. We employed the *TCGAbiolinks* function *GDCprepare* to obtain a Summarized Experiment object ^96^. We removed outlier samples with the *TCGAanalyze_Preprocessing* function of *TCGAbiolinks* using a Spearman correlation cutoff of 0.6. We normalized the datasets for GC-content ^97^ and library size using the *TCGAanalyze_Normalization* function of *TCGAbiolinks*. Lastly, we filtered the normalized RNA-Seq data for low counts across samples using the function *TCGAanalyze_Filtering* with a 0.20 cutoff for quantile filtering. For the TCGA datasets where normal samples were missing, we used the unified dataset that integrates the Genotype-Tissue Expression (GTEx) datasets ^52^ of healthy samples and the TCGA data, as provided by the *Recount2* protocol ^51^. We also employed this dataset as an additional source of information for the TCGA datasets with less than five normal samples (see Table S1). We used the *TCGAquery_Recount2* function of TCGAbiolinks ^98^ to query the GTEx and TCGA unified datasets. We carried out GC-content normalization and quantile filtering on the unified datasets, as described above.

Differential expression analyses have been carried out using *limma-voom* ^99^ as implemented within the *TCGAanalyze_DEA function* of *TCGAbiolinks* ^98^, along with *edgeR* to confirm the results, as we recently applied to another case study ^100^. We included in the design matrix conditions (tumor vs normal) and the TSS (Tissue Source Site; the center where the samples are collected) or the Plates (where available) as source of batch-effects to assess the robustness of the estimate of changes in expression with respect to different correction factors. In all our DEA analyses, we defined as a cutoff to retain significant DE genes a log fold change (logFC) >= 0.5 or <=-0.5, whereas a cutoff of 0.05 was used for the False Discovery Rate (FDR). We then retrieved the estimate logFC for ULK1 (ENSEMBL ID ENSG00000177169) and ULK2 (ENSG00000083290) in the different comparisons (**see Table S1**).

### Curation and analyses of missense mutations of ULK1 from TCGA

We retrieved mutations for ULK1 from each TCGA cancer study using the *MuTect2* pipeline ^101^ as implemented in the *TCGAbiolinks* function *GDCquery_Maf*. We retained missense mutations in the kinase domain of ULK1 for the structural analysis. For each mutation, we also collected the following information: i) potential of cancer driver mutation with *Fathmm* ^102^; ii) pathogenic potential with *PolyPhen* ^103^ and *SIFT* ^104^; iii) *REVEL* score ^105^; iv) interplay with post-translational modifications and functional short linear motifs using as a source of information *PhosphoSite* ^106^ and *ELM* ^107^, respectively; v) identification of the same mutation in *COSMIC* ^108^. We verified that the mutations under investigation were not found in *ExAC* as natural polymorphisms with high frequency in the healthy population ^109^. We also used *iSNO-AAPair* ^110^*, SNOSite* ^111^ and *NetPhos* ^112^ to predict *S*-nitrosylation or phosphorylation sites upon mutation to cysteine or phosphorylatable (serine, threonine and tyrosine) residues, respectively. We used the *NetPhos* predictor only for those mutations that were in solvent exposed sites upon analyzes of solvent accessibility of their sidechain with *NACCESS* (http://wolf.bms.umist.ac.uk/naccess/). Moreover, we verified that each mutation under investigation was the only one targeting the ULK1 gene in the sample where it was identified.

### Interactome of ULK1 and co-occurrence of mutations

We retrieved the experimentally known ULK1 interactors through the *Integrated Interaction Database* (IID) version 2018-05 ^75^. We then estimated the co-occurrence of mutations between ULK1 and each of these interactors, along with other ULKs kinases (i.e. ULK2, ULK3 and ULK4) with the *somaticInteractions* function of *maftools* R/Bioconductor package ^113^, which performs a pairwise Fisher’s Exact test to detect significant pairs of genes.

### Prediction of driver genes

We used the *oncodrive* function of *maftools* ^113^ to evaluate if ULK1 was predicted as driver gene in any of the cancer type under investigation. The function is based on the algorithm *oncodriveCLUST* ^73^.

### Free energy calculations

We employed the *FoldX* energy function ^114,115^ to perform in silico saturation mutagenesis. Calculations with this empirical energy function resulted in an average ΔΔG (differences in ΔG between mutant and wild-type variant) for each mutation over five independent runs performed using: i) the X-ray structure of ULK1 (PDB entry 5CI7, ^32^), ii) an ensemble of 20 representative conformations from the MD simulations with CHARMM22star or iii) with CHARMM27. The protocol is detailed in our previous publication ^46^. We also performed a literature-based curation of mutations for which the effects have been studied experimentally and use them as a control of the quality of our predictions.

### Molecular dynamics simulations

We carried out 1-μs molecular dynamics (MD) simulations for the human ULK1 kinase domains in explicit solvent using GROMACS software version 4.6 ^116^. We used as starting structure the PDB entry 5CI7 after in silico retro-mutation of the phospho-Thr 180 to Thr, to provide a model of the unphosphorylated variant of the domain. We used two protein different force fields CHARMM22* ^117^, CHARMM27 ^118^ in combination with the TIP3P water model ^119^ to evaluate the robustness of our results with respect to different physical models.

We used a dodecahedral box applying periodic boundary conditions and a concentration of NaCl of 150mM, neutralizing the charges of the system. The simulated system (protein+water) accounted for 83077 atoms. The system was prepared by different steps of minimization, solvent equilibration, thermalization and pressurization. We carried out productive MD simulations in the canonical ensemble at 300 K using velocity rescaling with a stochastic term ^120^. We applied the LINCS algorithm ^121^ to constrain the heavy atom bonds to use a time-step of 2 fs. We calculated long-range electrostatic interactions using the Particle-mesh Ewald (PME) summation scheme ^122^, whereas we truncated Van der Waals and short-range Coulomb interactions at 10 Å.

We verified the absence of artificial contacts between the periodic images of the protein in the simulations, which were always at a distance higher than 30 Å. We evaluated the quality of the conformational ensemble on 100 representative conformations equally spaced in time for each of the simulations using the machine-learning based approach implemented in *ResProx* ^123^ to predict the atomic resolution from structural ensembles of proteins. We obtained a predicted resolution of 1.53 +/− 0.30 and 1.66 +/− 0.12 Å for CHARMM22star and CHARMM27 simulations of ULK1 kinase domain, respectively. These values are very close to the resolution of the corresponding X-ray structure (1.74 Å), suggesting an overall quality of the MD-based ensemble.

We used Principal Component Analysis (PCA) of the covariance matrix of Cα atomic fluctuations to extract the principal motions from the MD simulations ^124^. We performed the PCA on a concatenated trajectory of the two MD simulations of ULK1 to compare them in the same essential subspace.

### CABS_flex ensembles of selected ULK1 mutant variants

For a selection of mutant variants of ULK1 (F14L, N96D, A101T, A125T, R137C, R137H and D138N) that have hub-behavior, we also collected conformational ensembles using the coarse-grained approach implemented in *CABS_flex* 2.0 ^125^.

We used as starting structures for *CABS_flex* calculations the models generated by *FoldX* during the mutational scan for each of these mutations. In particular, we selected the most representative models in terms of rotameric state of the mutated residue for each mutation. We then collected an ensemble of ten different conformations for each mutant variant to be used for the contact-based PSN analyses of hub residues described below, upon reconstruction of the corresponding full-atom models.

### Protein structure networks

We employed a contact-based Protein Structure Network (PSN) to the MD ensemble as implemented in *Pyinteraph* ^86^. We defined as hubs those residues of the network with at least three edges ^43^. We used the node inter-connectivity to calculate the connected components, which are clusters of connected residues in the graph. We selected 5 Å as the optimal cutoff for the contact-based PSN using the *PyInKnife* pipeline ^87^. The distance was estimated between the center of mass of the residue side chains. We removed spurious interactions during the simulations applying a persistence cutoff of 20% (i.e., each contact was included as an edge of the PSN only if occurring in 20% of the MD frames), as indicated in the original implementation of the method ^86^. We applied a variant of the depth-first search algorithm to identify the shortest path of communication. We defined the shortest path as the path in which the two residues were non-covalently connected by the smallest number of intermediate nodes.

We also calculated the persistence of salt bridges and hydrogen bonds with *PyInteraph* and the corresponding networks. For salt-bridges, all the distances between atom pairs belonging to charged moieties of two oppositely charged residues were calculated. The charged moieties were considered as interacting if at least one pair of atoms was found at a distance shorter than 4.5 Å. In the case of aspartate and glutamate residues, the atoms forming the carboxylic group were considered. The NH3- and the guanidinium groups were employed for lysine and arginine, respectively. We also verified the consistency of the results with a cutoff of 5 Å. We applied a persistence cutoff to filter interactions of 20% also for these networks.

## CONCLUSIONS

The assessment of the different effects that a mutation can exert on a protein explored in this study and the subsequent classification of the mutations can provide a useful complement to cancer genomics studies. For example, it allows to identify mutations that are likely to be ‘driver’ or ‘passenger’, along with to predict if the effect is triggered more by a destabilization of the protein product or a protein variant with impaired functionality. Moreover, our combined approach for mutation assessment could also benefit for the prioritization and selection of mutant variants for cellular experimental validation. Indeed, it can suggest how to select the proper readout for experimental validation. As an example, in a case where the mutation is predicted to be damaging for stability, experiments to estimate its cellular levels and half-life could be used, along with readouts to evaluate if the changes are due to proteasomal degradation or other clearance mechanisms. On the other side, if a mutation is predicted to result in a variant which is as stable as the wild-type, but the effect is more related to its function, experiments to evaluate its interactions in the cell with the biological partners, its regulation by PTMs or cellular assays to evaluate the effects on the pathways where interactors of the target protein are involved would be the most suitable choice. Moreover, we showed how the structural analyses used here benefit of the integration of bioinformatic tools to assess the changes in expression level of the target gene along with changes in other genes that can have compensatory effects, as we exemplified for ULK2. In addition, the extension of the analyses to the protein target interactome in terms of understanding co-occurrence of alterations and synergic effects that can arise from them allow a comprehensive view and to pinpoint interesting alterations at the molecular level. We here showed how our workflow can help in the study of a key kinase of the autophagy pathway, ULK1. We discovered that in the majority of the cases the gene expression levels are not altered or can be compensated by an up-regulation of the homologous kinase ULK2, whereas more than 30 different missense mutations altering the coding region of the gene have been identified. These mutations co-occur with mutations in ULK1 interactors fundamental for the upstream regulation of autophagy, suggesting an impairment of this process in cancer types such as uterine, stomach, skin, glioblastoma and colon cancers. Moreover, our study allowed to pinpoint that more than 50% of the mutations of ULK1 found in the cancer samples have an effect on protein stability, which is likely to have a more pronounced effect that the residual effect on protein activity, especially if it cannot be compensated by interactions with regulators of cellular ULK1 stability, which are also altered in the samples under investigation. We identified three mutations (S184F, D102N, and A28V) that predicted with only impact on kinase activity, either altering the functional dynamics of the protein or the capability to exert long range effects from distal site to the functional and catalytic regions. Future studies will be required to understand if these mutations have an inhibitory or activatory role on the kinase. The framework here applied could be more broadly extended to other targets of interest, as we recently started to apply, to help in the classification of mutational effects, along with prioritizing the variants for experimental validation and a specific biological readout.

## ACKNOWLEDGMENTS

The authors would like to thank Matteo Lambrughi and Matteo Tiberti for technical assistance and fruitful discussion. The results shown here are in part based upon data generated by the TCGA Research Network: https://www.cancer.gov/tcga. The project was supported by Danmarks Grundforskningsfond (DNRF125), Danish Council for Independent Research Project 1 (102517), and a Carlsberg Foundation Distinguished Fellowship (CF18-0314). Moreover, the project has been supported by a Netaji Subhash ICAR international fellowship, Govt. of India to MK to work in EP group. The calculations described in this paper were performed using the DeiC National Life Science Supercomputer Computerome at DTU (Denmark), and DECI-PRACE 14^th^ and 15^th^ HPC Grants for calculations on Archer (UK).

## Author contributions

Conceptualization: EP; Data Curation: MK, EP; Formal Analysis: MK, EP; Funding Acquisition: MK, EP; Investigation: MK, EP; Methodology: EP; Project Administration: EP; Resources: EP; Supervision: EP; Validation: MK, EP; Visualization: EP; Writing-Original Draft: EP; Writing-Review and Editing: MK, EP.

