## Supplementary TableS7 for "A Pan-Cancer assessment of alterations of the kinase domain of ULK1, an upstream regulator of autophagy"

**Table S7. Shortest paths of communication in the contact-based PSNs of CHARMM22star and CHARMM27 MD ensembles.** We used as target residues: i) residues important for activity (K46, E63, M62, T180 and K162); ii) residues of the DFG, HRD, and APE motifs; iii) central residues of the C-helix (55-65); iv) the residues of the catalytic and v) regulatory spines. No paths were found from the mutation sites to the regulatory spine or the APE motif in both the MD simulations.

| **mutation site- target site** | **class of target residues** | **path length (C22star)** | **average weight (C22star)** | **path length (C27)** | **average weight (C27)** |
| --- | --- | --- | --- | --- | --- |
| N96-K46 | Activity | 7 | 53 | 8 | 47.4 |
| A101-K46 | Activity | 9 | 69.4 | 10 | 58.7 |
| R137-K46 | Activity | 9 | 56.2 | 10 | 48.4 |
| D138-K46 | Activity | 7 | 63 | 8 | 53.8 |
| A28-K162 | Activity | 6 | 60.7 | 7 | 56.9 |
| N96-K162 | Activity | 3 | 49.4 | 3 | 40.8 |
| A101-K162 | Activity | 9 | 73.9 | 9 | 66.0 |
| R137-K162 | Activity | 9 | 60.7 | 9 | 54.5 |
| D138-K162 | Activity | 7 | 69 | 7 | 62.7 |
| A28-A44 | Catalytic spine | 4 | 62.1 | 5 | 59.2 |
| N96-A44 | Catalytic spine | 7 | 61.9 | 7 | 58.2 |
| A101-A44 | Catalytic spine | 9 | 76 | 9 | 68.2 |
| R137-A44 | Catalytic spine | 9 | 62.8 | 9 | 56.7 |
| D138-A44 | Catalytic spine | 7 | 71.9 | 7 | 65.6 |
| A28-L145 | Catalytic spine | 6 | 73.6 | 7 | 65.5 |
| N96-L145 | Catalytic spine | 4 | 86.7 | 4 | 76.9 |
| A101-L145 | Catalytic spine | 7 | 75.1 | 7 | 66.6 |
| D102-L145 | Catalytic spine | 9 | 76.8 | 9 | 66.0 |
| R137-L145 | Catalytic spine | 7 | 57.5 | 7 | 51.2 |
| D138-L145 | Catalytic spine | 5 | 68.5 | 5 | 62.0 |
| A28-V30 | Catalytic spine | 3 | 57.2 | 3 | 68.5 |
| N96-V30 | Catalytic spine | 8 | 49.8 | 9 | 47.2 |
| D138-V30 | Catalytic spine | 8 | 58.4 | 9 | 52.8 |
| N96-L146 | Catalytic spine | 4 | 33.3 | 4 | 52.7 |
| A101-I210 | Catalytic spine | 3 | 81.3 | 3 | 86.0 |
| D102-I210 | Catalytic spine | 5 | 81.7 | 5 | 75.0 |
| D138-I210 | Catalytic spine | 5 | 65.8 | 5 | 64.1 |
| R137-I210 | Catalytic spine | 7 | 55.7 | 7 | 52.7 |
| N96-L59 | C-helix | 10 | 65.9 | 10 | 60.7 |
| A101-L59 | C-helix | 6 | 53.5 | 6 | 52.2 |
| D102-L59 | C-helix | 8 | 61.7 | 8 | 55.6 |
| R137-L59 | C-helix | 3 | 30.1 | 3 | 31.9 |
| D138-L59 | C-helix | 3 | 29.4 | 3 | 31.9 |
| N96-L60 | C-helix | 8 | 50.4 | 9 | 46.8 |
| A101-L60 | C-helix | 10 | 65.5 | 9 | 46.8 |
| D138-L60 | C-helix | 8 | 59 | 9 | 52.4 |
| N96-D165 | DFG motif | 6 | 86.2 | 6 | 73.1 |
| D138-D165 | DFG motif | 3 | 51.5 | 3 | 56.6 |
| A28-D165 | DFG motif | 6 | 71.8 | 7 | 59.1 |
| D102-D165 | DFG motif | 7 | 74 | 7 | 65.5 |
| R137-D165 | DFG motif | 5 | 43.5 | 5 | 43.2 |
| A101-D165 | DFG motif | 5 | 69.9 | 5 | 66.3 |
| N96-D138 | HRD motif | 8 | 76.3 | 8 | 68.4 |
| A28-D138 | HRD motif | 8 | 30.3 | 9 | 48.7 |
| D102-D138 | HRD motif | 6 | 74.5 | 6 | 64.3 |
| A101-D138 | HRD motif | 4 | 69.6 | 4 | 64.6 |
| N96-R137 | HRD motif | 10 | 67.2 | 10 | 59.8 |
| D102-R137 | HRD motif | 8 | 63.4 | 8 | 54.5 |
