## Supplementary figures and images for "A Pan-Cancer assessment of alterations of the kinase domain of ULK1, an upstream regulator of autophagy"

### Figure S1

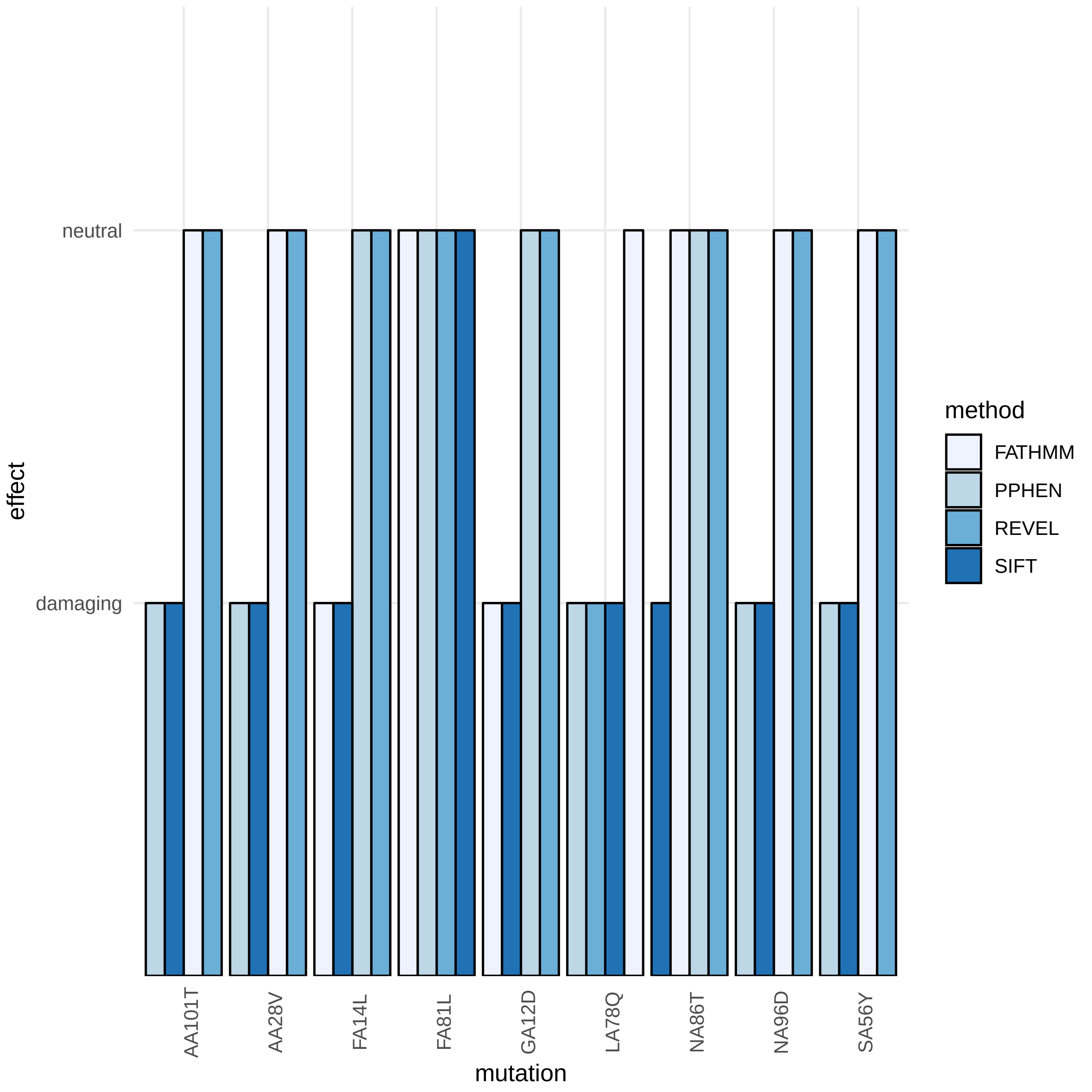

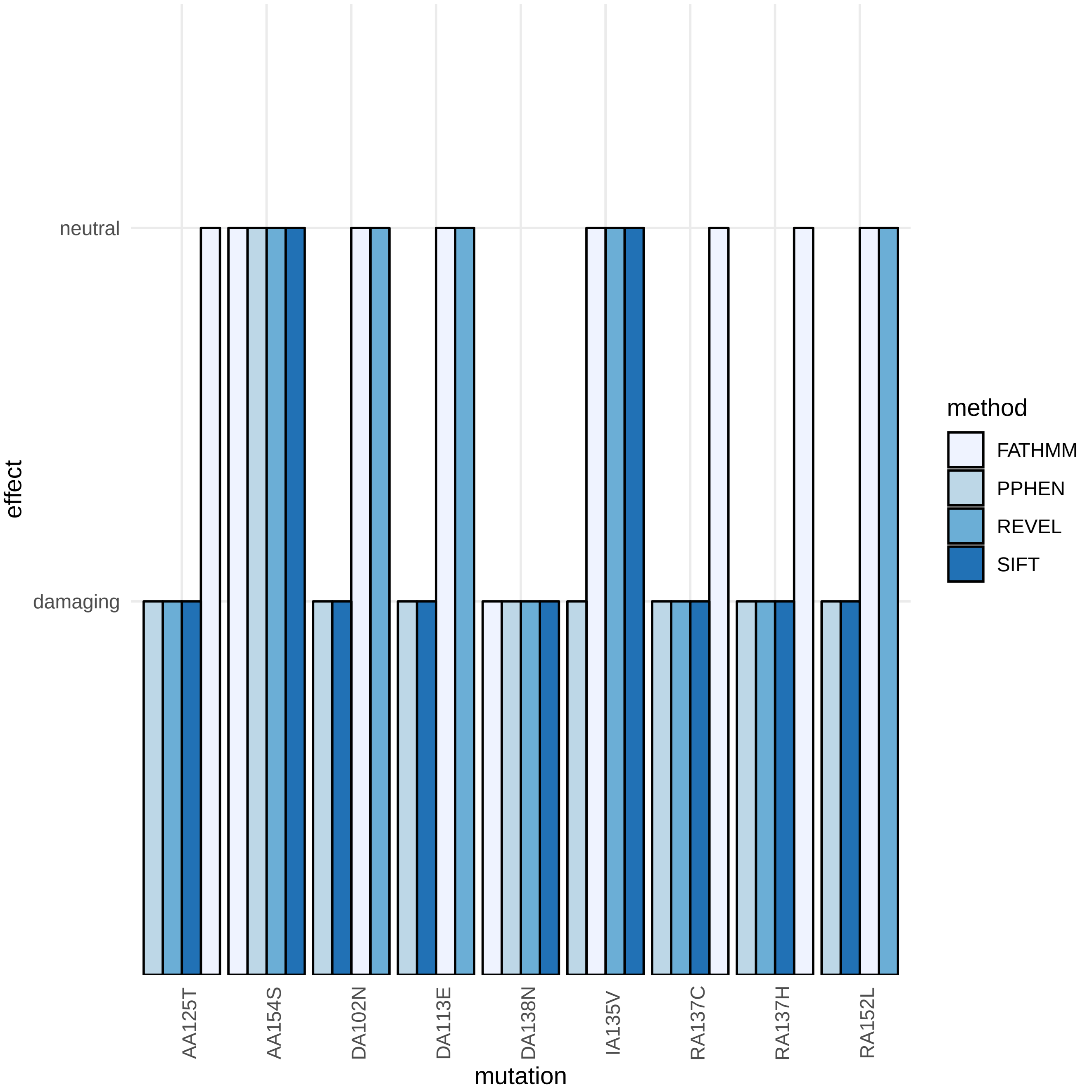

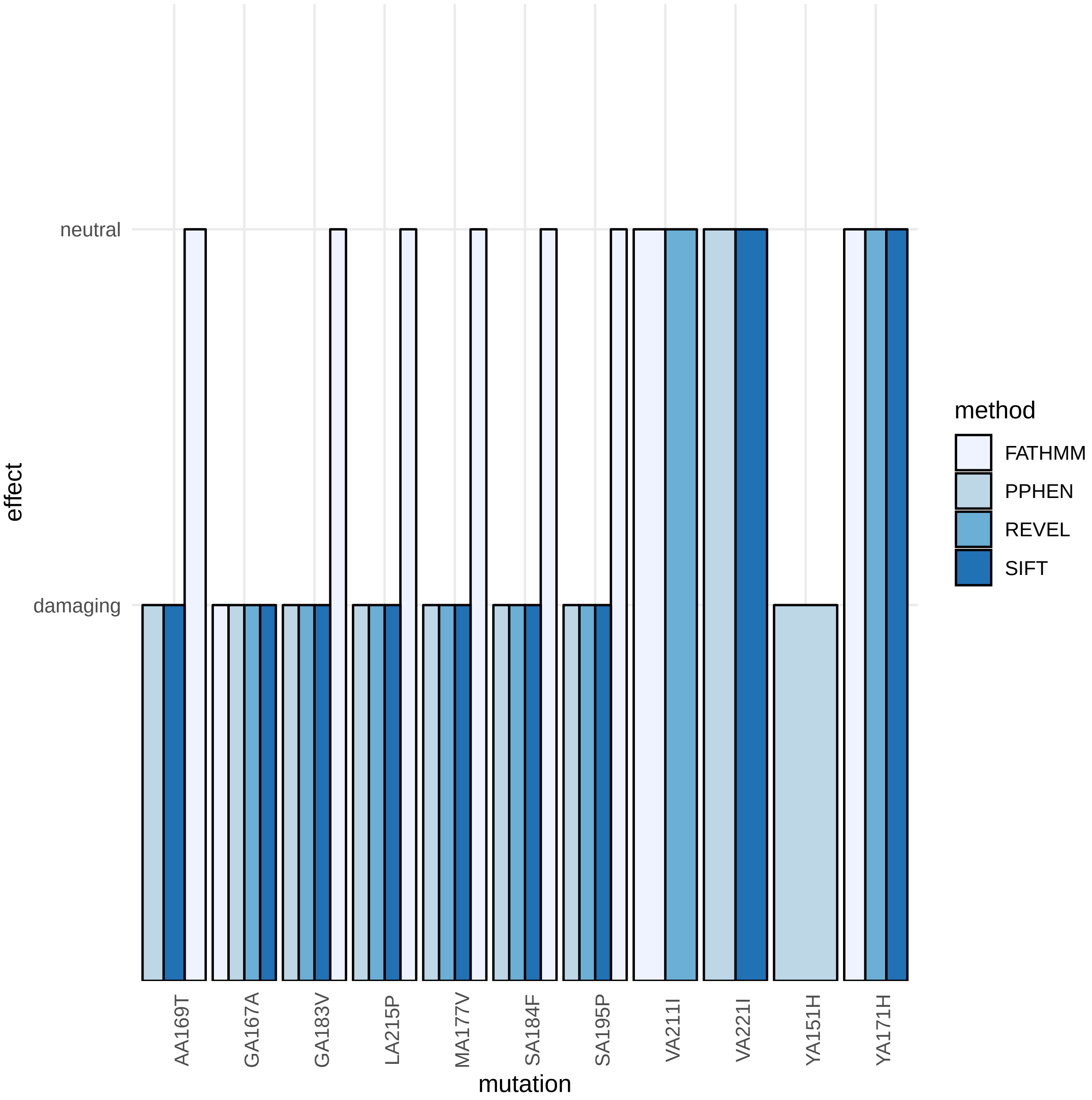
